# Differences in Hepatocellular Iron Metabolism Underlie Sexual Dimorphism in Hepatocyte Ferroptosis

**DOI:** 10.1101/2023.06.24.546395

**Authors:** Hui Tao, Hamid Y Dar, Cheng Tian, Somesh Banerjee, Evan S Glazer, Shanthi Srinivasan, Liqin Zhu, Roberto Pacifici, Peijian He

## Abstract

Males show higher incidence and severity than females in hepatic injury and many liver diseases, but the mechanisms are not well understood. Ferroptosis, an iron-mediated lipid peroxidation-dependent death, plays an important role in the pathogenesis of liver diseases. We determined whether hepatocyte ferroptosis displays gender difference, accounting for sexual dimorphism in liver diseases. Compared to female hepatocytes, male hepatocytes were much more vulnerable to ferroptosis by iron and pharmacological inducers including RSL3 and iFSP1. Male but not female hepatocytes exhibited significant increases in mitochondrial Fe^2+^ and mitochondrial ROS (mtROS) contents. Female hepatocytes showed a lower expression of iron importer transferrin receptor 1 (TfR1) and mitochondrial iron importer mitoferrin 1 (Mfrn1), but a higher expression of iron storage protein ferritin heavy chain 1 (FTH1). It is well known that TfR1 expression is positively correlated with ferroptosis. Herein, we showed that silencing FTH1 enhanced while knockdown of Mfrn1 decreased ferroptosis in HepG2 cells. Removing female hormones by ovariectomy (OVX) did not dampen but rather enhanced hepatocyte resistance to ferroptosis. Mechanistically, OVX potentiated the decrease in TfR1 and increase in FTH1 expression. OVX also increased FSP1 expression in ERK-dependent manner. Elevation in FSP1 suppressed mitochondrial Fe^2+^ accumulation and mtROS production, constituting a novel mechanism of FSP1-mediated inhibition of ferroptosis. In conclusion, differences in hepatocellular iron handling between male and female account, at least in part, for sexual dimorphism in induced ferroptosis of the hepatocytes.

## 1. Introduction

The liver is the hub of numerous physiological processes including nutrient metabolism, immune support, cholesterol homeostasis, blood volume regulation, and the breakdown of xenobiotic compounds [1]. Liver diseases are often inflammatory in nature and may be resulted from viral infection, drug overdose, overnutrition, and alcohol abuse [1]. The absolute number of chronic liver disease cases is estimated at 1.5 billion worldwide, including non-alcoholic fatty liver disease (NAFLD, 59%), hepatitis B (29%), hepatitis C (9%), and alcoholic liver disease (ALD, 2%) [2]. The above-mentioned factors cause endoplasmic reticulum stress, oxidative stress, and mitochondrial dysfunction in hepatic cells particularly the hepatocytes [3–5]. Unresolved stress further leads to hepatic cell death and liver injury that in turn triggers liver inflammation, which over time causes liver fibrosis, cirrhosis, and cancer [6]. Interestingly, the prevalence and severity of liver diseases display gender difference. Compared to women, men are more likely to develop chronic liver disease with two to three folds higher likelihood of having cirrhosis and hepatocellular carcinoma [7], the most common type of primary liver cancer that accounts for 80-90% of all liver cancers.

Iron is vital for hemoglobin production and cellular metabolism. However, excess iron is hazardous by causing oxidative damage of intracellular molecules (DNA/lipid/protein) via the Fenton reaction [8]. Elevated liver iron loads are often seen in patients with hereditary hemochromatosis, ALD, and viral hepatitis [9]. A mild-to-moderate increase of liver iron also occurs in one-third of patients with NAFLD [10]. Hepatic iron overload promotes the pathogenesis and progression of liver diseases by inducing liver injury, inflammation, and fibrosis, ultimately leading to cirrhosis and liver cancer. Ferroptosis, a recently defined iron-mediated lipid peroxidation-dependent oxidative cell death [11], plays an important role in the pathogenesis of many liver diseases including acetaminophen-induced acute liver failure, hepatic ischemia-reperfusion injury, viral hepatitis, and steatohepatitis [12]. Ferroptosis is featured by disturbed iron homeostasis and impaired antioxidant defense [13]. Iron catalyzes the peroxidation of poly-unsaturated fatty acids, while antioxidant defect or deficit results in uncontrollable propagation of lipid peroxidation, ultimately leading to ferroptosis. Glutathione peroxidase 4 (GPX4) and ferroptosis suppressing protein 1 (FSP1), which is formerly known as apoptosis-inducing factor mitochondria associated protein 2 (AIFM2), are the two key antioxidant enzymes that protect against ferroptosis [11, 14, 15]. GPX4 utilizes glutathione as a cofactor to protect against lipid peroxidation [11], and FSP1 catalyzes the production of reduced coenzyme Q10 (CoQH_2_) [15, 16], which traps lipid peroxyl radicals.

A recent study showed that renal epithelial cell ferroptosis displays gender difference accounting for sexual dimorphism in kidney injury [17]. We have recently shown that iron-loaded hepatocytes undergo ferroptosis because of iron-catalyzed overproduction of mitochondrial reactive oxygen species (mtROS) [18]. Intriguingly, sexual dimorphism in iron metabolism has been well noted in humans and in experimental rodents [19–21]. Hepatocytes, which compromise approximately 80% of the liver mass, play an important role in systemic iron homeostasis via the deposition of excessive body iron and production of hepcidin [22], a master regulator of iron metabolism. We wondered whether the induced ferroptosis of hepatocytes manifests a sex difference, and whether the potential differences in iron metabolism are the underlying mechanism. Hepatocytes primarily take up iron in the form of transferrin-bound iron via transferrin receptor 1 (TfR1) [23], and uptake non-transferrin-bound iron (NTBI) via Zrt- and Irt-like (ZIP) protein ZIP14 when iron is in excess [24]. Divalent metal transporter 1 (DMT1) then transports iron out of the endosomes for intracellular utilization [25]. A key site of iron utilization is the mitochondria, which is highly enriched in hepatocytes to meet the need of metabolism. The mitochondrial iron transport process is not fully clear but potentially involves DMT1 [26], mitoferrin 1 and 2 (Mfrn1/2) [27], and mitochondrial Ca^2+^ uniporter (MCU) [28]. Excess iron is then stored in the ferritin complexes in the cytosol to limit free iron mediated oxidative damage to cellular contents [29].

In this study, by utilizing multiple different inducers of ferroptosis, we determined whether ferroptosis of the hepatocytes display gender difference by using primary hepatocytes from male and female mice as well as mice with ovariectomy (OVX). We have further analyzed the expression profiles of genes associated with hepatocellular iron metabolism and antioxidant defense, which protects against ferroptosis, to identify the underlying mechanisms. We demonstrated sex dimorphism in the ferroptosis of hepatocytes, which is at least in part due to differences in hepatocellular iron handling. Intriguingly, ovariectomy did not impair but strengthened the resistance of female hepatocytes to ferroptosis.

## Materials and methods

### Reagents

The materials and reagents used in the current study are listed in **Supplementary Table 1**.

### Animals

Male and female C57BL/6 mice were purchased from the Jackson Laboratory. Female mice were either sham-operated or ovariectomized by the dorsal approach under general anesthesia at 8 weeks of age [30], and then sacrificed two weeks later. All animal procedures were approved by the Institutional Animal Care and Use Committee of Emory University.

### Hepatocyte culture

Primary mouse hepatocytes (P-Hepa) were isolated from 8-10 weeks old male and female mice as previously described [18, 31]. Briefly, mouse liver was pre-perfused with Leffert’s buffer (HEPES 10 mM, KCl 3 mM, NaCl 130 mM, NaH_2_PO_4_ 1 mM, glucose 10 mM, pH 7.4) containing EGTA, followed by perfusion with Liberase that consists of collagenase I and II. Dissociated hepatocytes were purified on a Percoll gradient, and cell viability is determined by staining with 0.02% trypan blue. Only isolations that have a viability of >90% were used. Male and female P-Hepa were seeded in collagen-coated 12-well plates at the density of 2 x 10^5^ cells per well for 3 h in William’s E medium supplemented with 5% FBS, 1x P/S, 10 mM HEPES, 1x insulin and 40 ng/ml dexamethasone. P-Hepa were then cultured overnight in fresh medium containing 5% FBS, 1x P/S and 10 mM HEPES prior to any treatments. HepG2 cells, a human hepatoblastoma cell line, were cultured in RPMI medium that is supplemented with 10% FBS and 1x P/S. AML12 cells are mouse hepatocytes isolated from a transgenic mouse that expresses human transforming growth factor α (TGF-α). AML12 cells were cultured in DMEM/F12 medium supplemented with 10% FBS, 1x P/S, 1x ITS, and 40 ng/ml dexamethasone. All cells were cultured in a 5% CO_2_ humidified incubator at 37°C.

### Lentivirus infection

Plasmid constructs pLKO.1/shMfrn1 (TRCN0000044010) and pLKO.1/shFTH1 (TRCN0000029432) were purchased from Sigma-Aldrich, and V5-tagged FSP1 (V5-FSP1) in the pLX304 backbone was purchased from DNASU plasmid repository. Lentiviral particles were prepared in 293T cells as previously described [32], and then utilized for infection of HepG2 cells in the presence of 5 μg/ml polybrene. Stable expression of shMfrn1, shFTH1 or the scrambled control shRNA was selected for three passages with 5 μg/ml puromycin, and stable expression of V5-FSP1 was selected with 5 μg/ml blasticidin. Selection antibiotics were excluded when cells were seeded for experiments.

### Immunohistochemistry

The de-identified human liver samples of cancer patients, including five men and four women, were obtained under a protocol approved by the Institutional Review Boards at both St. Jude Children’s Research Hospital and The University of Tennessee Health Science Center. The basic information of the involved subjects is shown in **Supplementary Table 2**. Formalin-fixed paraffin-embedded sections (4 µm) of liver sections were prepared and subject to standard immunohistochemistry with anti-Mfrn1 antibody at 1:250 dilution. Images were obtained to show the expression of Mfrn1 in normal-looking area of the liver.

### Quantitative RT-PCR analysis

Total RNA was extracted from mouse livers and cultured cells with the RNeasy Mini Kit. Three μg of total RNA was used for first strand cDNA synthesis using the High-Capacity cDNA Reverse Transcription Kit according to the manufacturer’s instruction. Quantitative PCR was performed with *Power* SRBR Green PCR Master Mix on QuantStudio 3 real-time PCR system. PCR primer sequences are listed in **Supplementary Table 3**.

### Western blot analysis

Male and female mouse livers were homogenized in 1 × lysis buffer containing 20 mM Tris-HCl (pH 7.5), 150 mM NaCl, 1 mM β-glycerophosphate, 2.5 mM sodium pyrophosphate, 1 mM Na_2_-EDTA, 1 mM EGTA, 1 mM Na_3_VO_4_, 1 μg/ml leupeptin, 1% Triton X-100, and protease/phosphatase inhibitors [18]. Cultured cells were lysed in the same lysis buffer. The crude lysates were then sonicated for 2 × 15 s and spun at 14,000 *g* for 15 min. The protein concentration of the resulted supernatant was determined by Bicinchoninic Acid Protein Assay. Protein lysates were boiled at 95°C for 10 min in 1× Laemmli buffer, separated on SDS-PAGE gel, and transferred to nitrocellulose membrane for Western blotting with the corresponding antibodies listed in Supplementary Table 1. Densitometric analysis was performed by using the ImageJ software.

### Propidium iodide (PI) staining

Staining of live cells with PI was as previously described [18]. Briefly, treated hepatocytes were incubated for 30 min with 2 μg/ml PI in the culture medium. Cells were washed with PBS, followed by fluorescent imaging by the Olympus IX83 imaging system. The numbers of PI^+^ cells and total numbers of cells *per* field were counted, from which the percentage of PI^+^ cell death was calculated. Please note that many P-Hepa have two nuclei, and thus one necrotic cell may show two red puncta from the PI staining.

### Detection of mitochondrial ROS (mtROS) and Fe^2+^ contents

P-Hepa and HepG2 cells grown on 4-well Lab-Tek II chambered coverglass were incubated with 5 μM Mito-FerroGreen and/or 5 μM MitoSOX in culture medium for the detection of mitochondrial Fe^2+^ and mtROS, respectively. Thirty minutes after incubation, cells were washed two times with HBSS buffer and subject to confocal imaging using the Olympus FV1000 microscope. Fluorescence intensity was quantified using the ImageJ software, and the relative changes in percentage are presented.

### Lipid peroxidation assay

Cultured hepatocytes were incubated for 30 min with 2 μM BODIPY® 581/591 C11 (Image-iT Lipid Peroxidation Kit). After washing with PBS, cells were imaged at the emission of ∼590 nm (red) and ∼510 nm (green). A shift from red to green fluorescence signal suggests lipid oxidation.

### Statistical analysis

Data were analyzed with the GraphPad Prism software (GraphPad Software, La Jolla, CA, USA) and are presented as mean ± SEM. Statistical significance was determined by unpaired t-test, with a value of *P* < 0.05 was considered significant.

## Results

### Male hepatocytes are more susceptible to ferroptosis induction than female hepatocytes

We have recently shown that iron induces positive staining for PI in cultured P-Hepa, and this PI positivity was abolished by ferrostatin-1, a lipid ROS scavenger and inhibitor of ferroptosis, thereby indicating that acute iron loading in P-Hepa induces ferroptosis [18]. By continuing using this primary culture model and PI staining, the vulnerability of male and female hepatocytes to ferric ammonium citrate (FAC) induced ferroptosis was assessed. We observed that treatment by 100 μM FAC for 24 h led to a markedly higher rate of ferroptosis in male hepatocytes than in female hepatocytes as indicated by the percentage of PI^+^ cells (32.3±1.5 *vs* 6.4±0.6) (**Fig. 1A**). The difference was further confirmed by treating male and female hepatocytes with a lower (20 μM) and higher (200 μM) concentration of FAC for a shortened duration (4 h). In response to the lower concentration of FAC, 7.4% of male hepatocytes underwent ferroptosis whereas only 0.7% female cells were PI positive (**Fig. 1B**). Much higher numbers (30.5%) of male hepatocytes died from the treatment of 200 μM FAC, but there remained only 0.9% female hepatocytes positive for PI (**Fig. 1B**). These data demonstrate that, compared to female, male hepatocytes are much more sensitive to iron excess induced ferroptosis, at least in the cell culture model.

**Figure 1.**
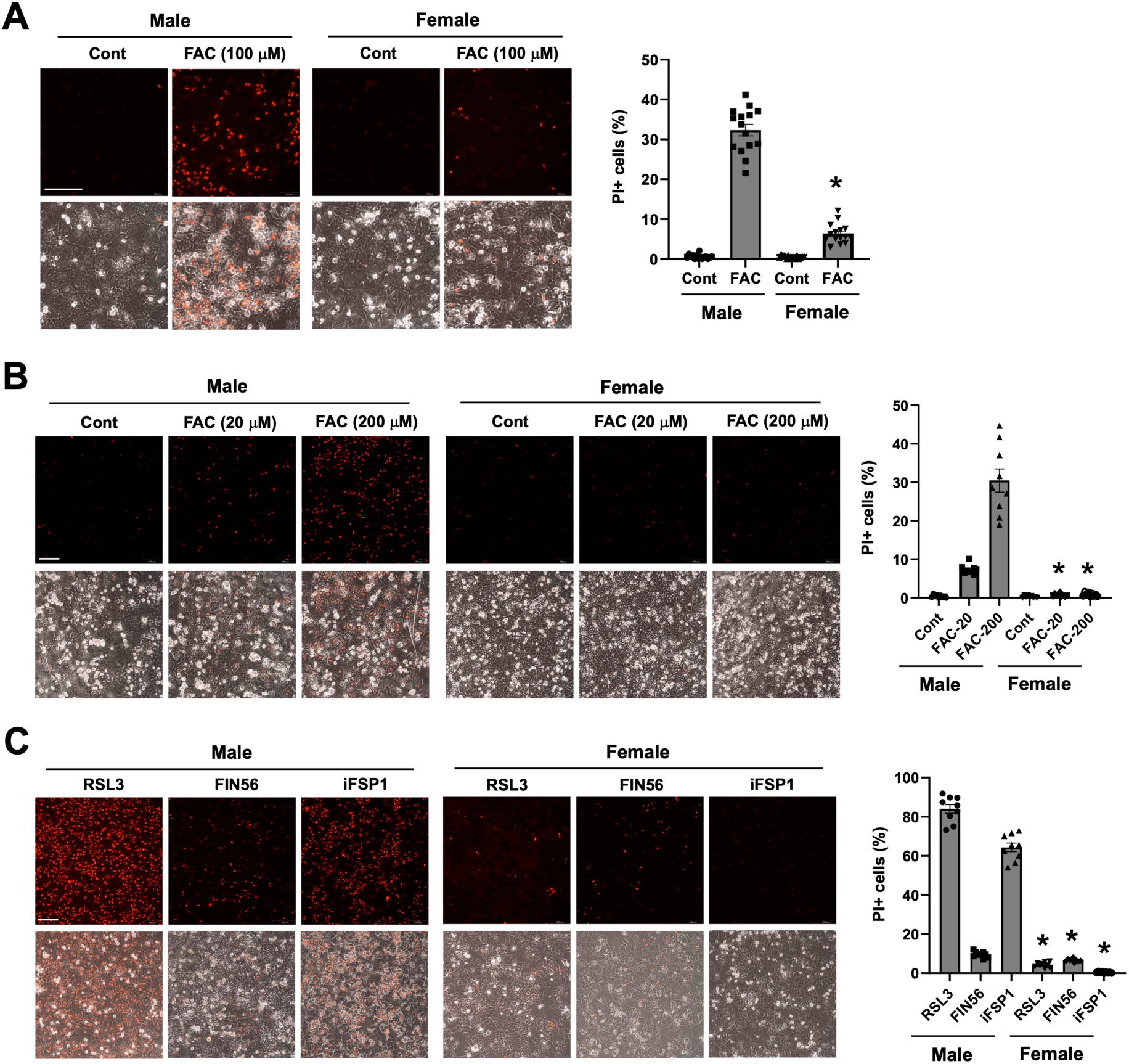
Differential vulnerability of male and female hepatocytes to ferroptosis. (**A**) Primary cultured mouse hepatocytes (P-Hepa) of male and female were treated with 100 μM FAC for 24 h and then stained for propidium iodide (PI). (**B**) P-Hepa were treated with 20 or 200 μM FAC for 4 h, followed by PI staining. (**C**) P-Hepa were treated with 100 nM RSL3 for 4 h, 2 μM FIN56 or 10 μM iFSP1 for 24 h prior to PI staining. The percentages of PI^+^ cells over the total number of cells were quantified and are shown next to the representative images. Data are mean ± SEM (n = 15 fields of 3 independent experiments for **A**, and 9 fields for **B** and **C**). *, *P* < 0.01 compared to the corresponding treatment conditions in male. Bars, 200 μm.

Multiple ferroptosis inducers including RSL3 [33], Ferroptosis inducer-56 (FN56) [34], and iFSP1 [35], have been used in studying ferroptosis induction of many cell species. It was shown that RSL3 directly inactivates GPX4, while FIN56 induces autophagy-dependent degradation of GPX4 [34]. Unlike RSL3 and FIN56, iFSP1 is a potent inhibitor of FSP1, which inhibits ferroptosis by catalyzing the production of antioxidant CoQH_2_ [35]. RSL3 as a strong inducer of ferroptosis was used at 100 nM for a 2-h treatment, while FIN56 and IFSP1 were used at micromolar levels for 24 h. Importantly, RSL3, FIN56 and iFSP1 all induced higher rates of ferroptosis in male than female hepatocytes, although the difference was small with FIN56 treatment (**Fig. 1C**). The gender differences in the magnitude of ferroptosis in responses to RSL3 and FIN56 may be explained by their different mechanisms of action. RSL3 induces overwhelming mtROS production and impaired mitochondrial membrane potential [36], while FIN56 does not perturb mitochondrial integrity [34]. Intriguingly, we observed that RSL3 and iFSP1, but not FIN56, induced marked increases in mitochondrial Fe^2+^ content and mtROS production in HepG2 cells, a hepatoblastoma cell line (**Fig. S1**). This finding led us suspect whether mitochondrial iron status and mtROS production are different between male and female hepatocytes in response to FAC and pharmacological inducers. Indeed, in response to FAC treatment, male hepatocytes exhibited a strong increase in the abundance of mitochondrial Fe^2+^ and mtROS, but corresponding changes were modest in female (**Fig. 2A**). A similar difference was seen between male and female hepatocytes in response to RSL3 and iFSP1, but FIN56 failed to induce a significant increase in mitochondrial iron or mtROS content in either sex (**Fig. 2B**). These findings demonstrate that male hepatocytes are more susceptible to ferroptosis induction, and the vulnerability of male hepatocytes to ferroptosis is likely associated with mitochondrial iron status and the capacity of mtROS production.

**Figure 2.**
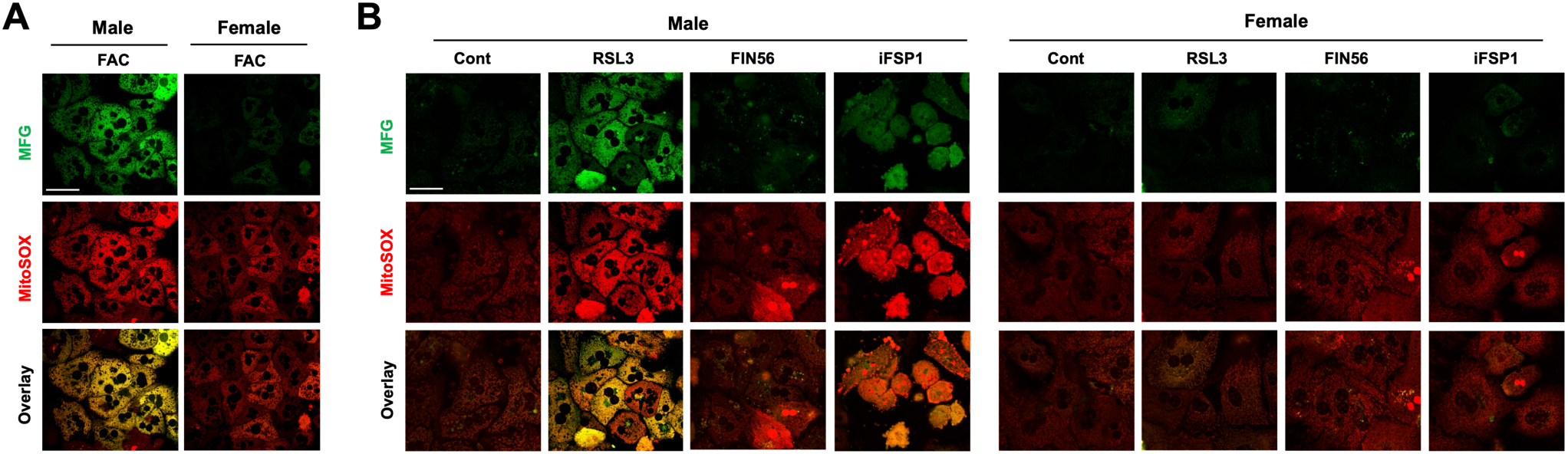
Sex difference in mitochondrial iron loading and mtROS production by ferroptosis inducers. (**A**) Male and female P-Hepa were treated with 100 μM FAC for 4 h, followed by staining for mitochondrial Fe^2+^ with 5 μM Mito-FerroGreen (MFG) and mtROS with 5 μM MitoSOX. (**B**) P-Hepa were treated with 100 nM RSL3 for 4 h, 2 μM FIN56 or 10 μM iFSP1 for 24 h prior to staining with MFG or MitoSOX. Bars, 50 μm.

### Differential expression in iron metabolism genes between male and female hepatocytes

We then interrogated whether hepatocellular iron metabolism is different between male and female hepatocytes. These hepatocellular iron metabolism related genes include TfR1 and ZIP14, ferritin heavy chain 1 (FTH1), and DMT1(+/- iron-responsive element, IRE), Mfrn1/2 and MCU. The expression of GPX4 and FSP1 was also determined given their critical roles in protecting against ferroptosis. We found that TfR1 and Mfrn1 transcript levels were significantly lower in female than in male livers, whereas the expression of DMT1(-IRE) and MCU was higher in female (**Fig. 3A**). There was not a sex difference in the mRNA expression of GPX4 or FSP1. At the protein level, TfR1 expression was consistently lower in female livers, but FTH1 expression was significantly higher in female (**Fig. 3B**). Again, we did not see a difference in the protein expression of GPX4 between male and female. The expression of nuclear factor E2-related factor 2 (NRF2), a key transcription factor that drives the antioxidant protection pathways, was also determined but showed no significant difference. The protein expression of Mfrn1 and FSP1 was not determined as we couldn’t obtain reliable antibodies against mouse Mfrn1 or FSP1 protein.

**Figure 3.**
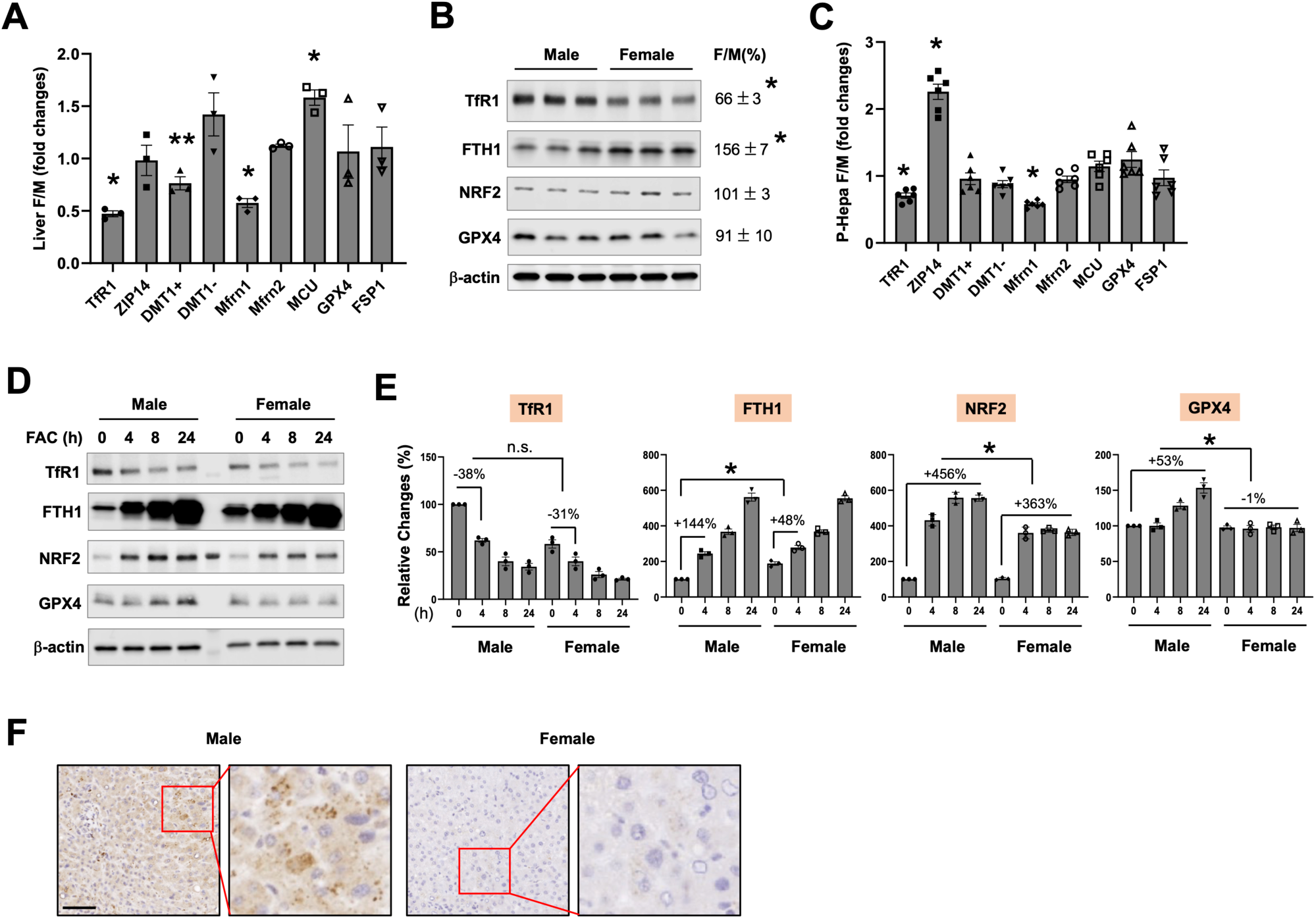
Expression profiling of genes associated with hepatocellular iron metabolism and antioxidant defense. The mRNA (**A**, **C**) and protein (**B**, **D**) expression of iron metabolism and antioxidant genes in the liver (**A**, **B**) and cultured P-Hepa (**C**, **D**) of male (M) and female (F) mice. DMT1+, DMT1 with IRE; and DMT1-, DMT1 without IRE. (**E**) Quantification of the relative changes in the protein expression of the determined genes in P-Hepa treated by 100 μM FAC for the indicated times. (**F**) Representative images showing Mfrn1 expression in the livers of male and female human subjects. Data are mean ± SEM (n = 3 for **A**, **B**, **D** and **E;** n = 6 for **C**). *, *P* < 0.01; and **, *P* < 0.05. n.s., not significant. Bar, 100 μm.

Consistent with findings in the liver, the transcript levels of TfR1 and Mfrn1 were significantly lower in female hepatocytes in *in vitro* (**Fig. 3C**). A lower TfR1 and higher FTH1 protein level was also found in female P-Hepa (**Fig. 3D and 3E**). FAC induced reduction in TfR1 expression was not significantly different between male and female at any of the examined time points. In contrast, FAC-induced increases in FTH1 expression were significantly smaller in female hepatocytes (**Fig. 3E**). The induction of NRF2 was also less robust in female, and GPX4 induction was only seen in male hepatocytes (**Fig. 3E**). We have validated an anti-human Mfrn1 antibody by using exogenously expressed human Mfrn1 and utilized this antibody to compare Mfrn1 expression in human livers. Consistent with Mfrn1 mRNA expression in mice, Mfrn1 expression in women’s liver was dramatically lower than that in men (**Fig. 3F**). This difference of Mfrn1 expression between men and women was consistently seen in all the currently analyzed human liver samples (**Fig. S2)**. These results demonstrate that male and female hepatocytes manifest significant differences in baseline expression of iron metabolism genes including TfR1, FTH1 and Mfrn1, but not antioxidant genes including GPX4, FSP1 and NRF2. Differential induction of GPX4 and NRF2 in response to FAC treatment more likely indicates different levels of oxidative damage caused to male and female hepatocytes.

### FTH1 and Mfrn1 regulate the sensitivity of hepatocytes to ferroptosis

The above data suggest a lower expression of TfR1 and Mfrn1 but a higher expression of FTH1 probably accounts, at least in part, for the lower vulnerability of female hepatocytes to ferroptosis. It was well documented that TfR1 expression level is positively correlated with the likelihood of developing ferroptosis [37]. In contrast, the role and mechanism of FTH1 (a major iron storage protein) and Mfrn1 (a mitochondrial iron importer) in the regulation of ferroptosis are less clear. Herein, we determined whether manipulating the expression of Mfrn1 and FTH1 modulates hepatocyte’s sensitivity to ferroptosis by using HepG2 cells. To this end, HepG2 cells with the knockdown (KD) of FTH1 or Mfrn1 were generated. The acquired HepG2 cell stains displayed 91% reduction of FTH1 expression (**Fig. 4A**) and a 51% decrease in Mfrn1 (**Fig. 4B**). We have also determined the effects of FTH1 or Mfrn1 deficiency on the expression of other iron metabolism genes as well as GPX4 and FSP1. Surprisingly, KD of FTH1 did not significantly alter the expression of TfR1, indicating that deficiency in an iron storage protein did not lead to cytosolic accumulation of labile iron, which would otherwise have led to TfR1 reduction to stop continuous iron import into the cells. Mfrn1 expression was slight decreased by KD of FTH1. There was not a significant change in the expression of GPX4, but a 37% increase in FSP1 was observed (**Fig. 4A**). In contrast to the effects by KD of FTH1, silencing Mfrn1 significantly decreased TfR1 and FTH1 expression by 56% and 73%, respectively. KD of Mfrn1 did not alter FSP1 expression but led to a 54% increase in GPX4 expression (**Fig. 4B**).

**Figure 4.**
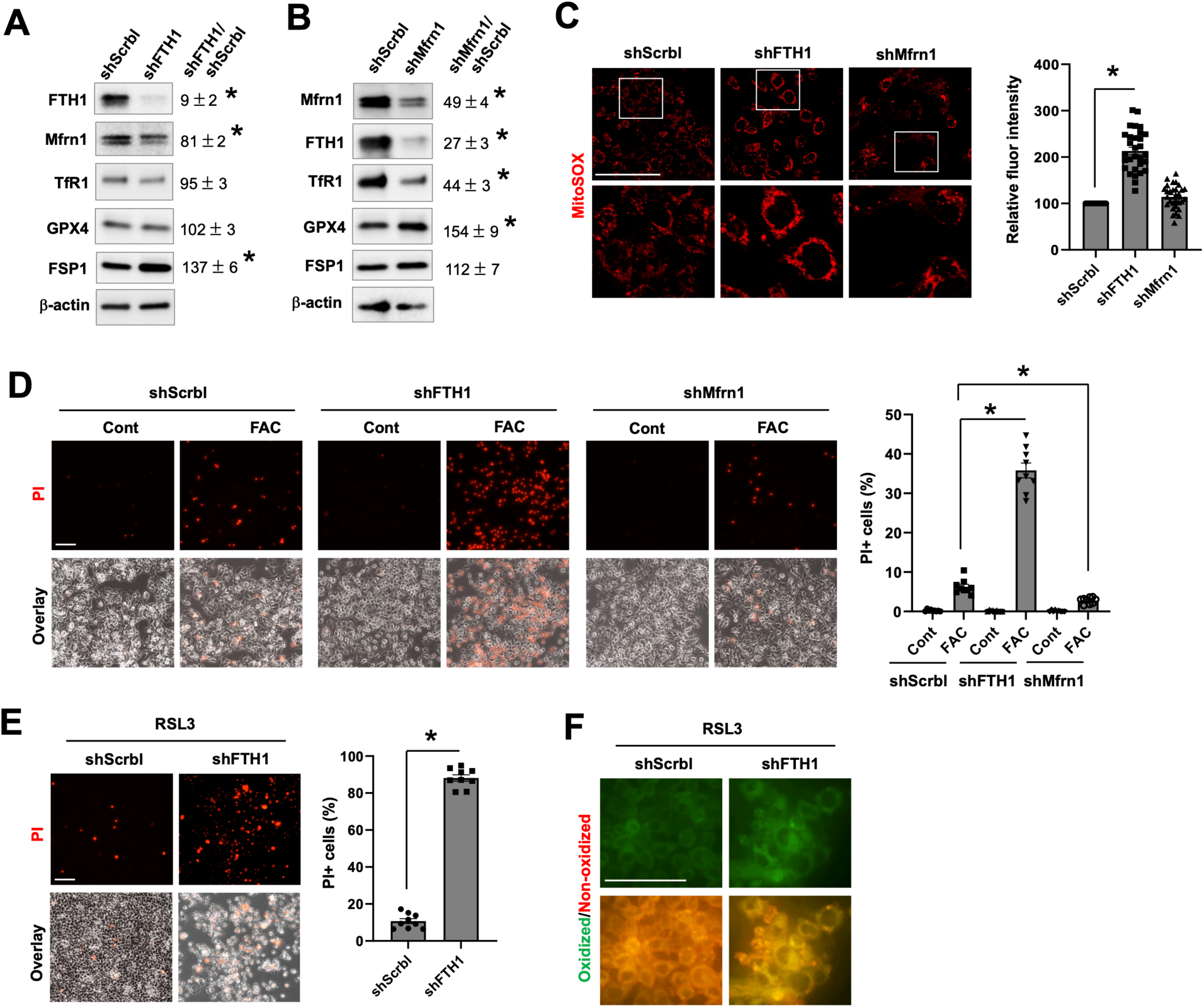
FTH1 and Mfrn1 regulate the sensitivity of hepatocyte to ferroptosis. The effects of the knockdown of FTH1 (shFTH1, **A**) and Mfrn1 (shMfrn1, **B**) on the protein expression of genes associated with iron metabolism and antioxidant defense by comparing to the scrambled control (shScrbl) in HepG2 cells. Next to the blots are the relative changes shown as mean ± SEM (n = 3). (**C**) Staining for baseline mtROS with MitoSOX in HepG2/shFTH1, HepG2/shMfrn1 and control shScrbl cells. The fluorescence intensity of MitoSOX staining was quantified and is presented as mean ± SEM (n = 30 cells of 3 independent experiments). (**D**) HepG2 cells with the knockdown of FTH1, Mfrn1 or the scrambled control were treated with 100 μM FAC for 24 h, and ferroptotic cells was stained by PI. (**E**) HepG2/shFTH1 and control shScrbl cells were treated with 100 nM RSL3 for 4 h, followed by PI staining. The percentages of PI^+^ cells (**D** and **E**) were quantified, and the data shown are mean ± SEM (n = 9 fields of 3 independent experiments). (**F**) Lipid peroxidation was determined by staining with BODIPY C11 in RSL3 treated shFTH1 and control cells, with the green fluorescence denoting lipid oxidation. *, *P* < 0.01. Bars, 100 μm.

Compared to the control cells, HepG2 cells with the KD of FTH1 displayed a significantly increased mtROS production, but deficiency in Mfrn1 did not change mtROS content at the baseline (**Fig. 4C**). Importantly, FTH1-deficient HepG2 cells were highly sensitive to FAC-induced ferroptosis, while Mfrn1 deficiency caused a slight but significant decrease in FAC-dependent ferroptosis (**Fig. 4D**). Deficiency in FTH1 also sensitized HepG2 cells to RSL3-induced ferroptosis (**Fig. 4E**), consistent with the significantly increased lipid peroxidation (**Fig. 4F**). We also determined the effect of Mfrn1 deficiency on RSL3- mediated ferroptosis but did not find a significant difference from the control cells (data not shown). An explanation for that Mfrn1-deficient cells were relative resistant to FAC, but not to RSL3, could be due to their differential effects on mitochondrial iron state and/or mtROS production. Collectively, our current findings demonstrate that hepatocytes with abundant ferritin are more protected from ferroptosis by different inducers, while low in Mfrn1 expression does not warrant a better resistance against ferroptosis but depends on the context.

### OVX enhances the defense of female hepatocytes against ferroptosis

Much of the protective nature in female attributes to female hormone-mediated signaling. We wondered whether this is true for the lower vulnerability of female hepatocytes to ferroptosis. To this end, OVX mice were generated, and OVX hepatocytes were prepared to be compared those of sham-female and male mice. Consistent with our earlier data, female hepatocytes were less susceptible to FAC-induced ferroptosis than male cells (**Fig. 5A**). Yet, to our surprise, OVX hepatocytes displayed increased resistance to FAC-induced ferroptosis compared to the sham-female control. Hepatocytes are an important site of the metabolism of macronutrients including glucose, sucrose, fatty acids, and amino acids [38]. Cellular metabolism of these nutrients involves the mitochondria, which plays a crucial role in ferroptosis. We showed that addition of glucose, fructose, palmitic acid (the most common long-chain fatty acid) or amino acids in the culture medium along with FAC all resulted in massive increase in the number of PI^+^ cells than FAC alone (**Fig. 5B**). The numbers of PI^+^ cells were significantly fewer for all conditions in sham-female hepatocytes and even less in OVX hepatocytes (**Fig. 5B**). We confirmed whether the different rates of PI^+^ cell death among male, sham-female, and OVX hepatocytes are associated with different levels of lipid peroxidation, a hallmark feature of ferroptosis. The FAC and palmitic acid cotreatment condition was selected because prominent differences in the rate of ferroptosis were seen among male, female and OVX hepatocytes. As expected, the co-treatment led to the highest level of lipid oxidation in male hepatocytes and least in OVX hepatocytes (**Fig. 5C**). These findings demonstrate that removal of female hormones enhances the resistance of female hepatocytes to ferroptosis.

**Figure 5.**
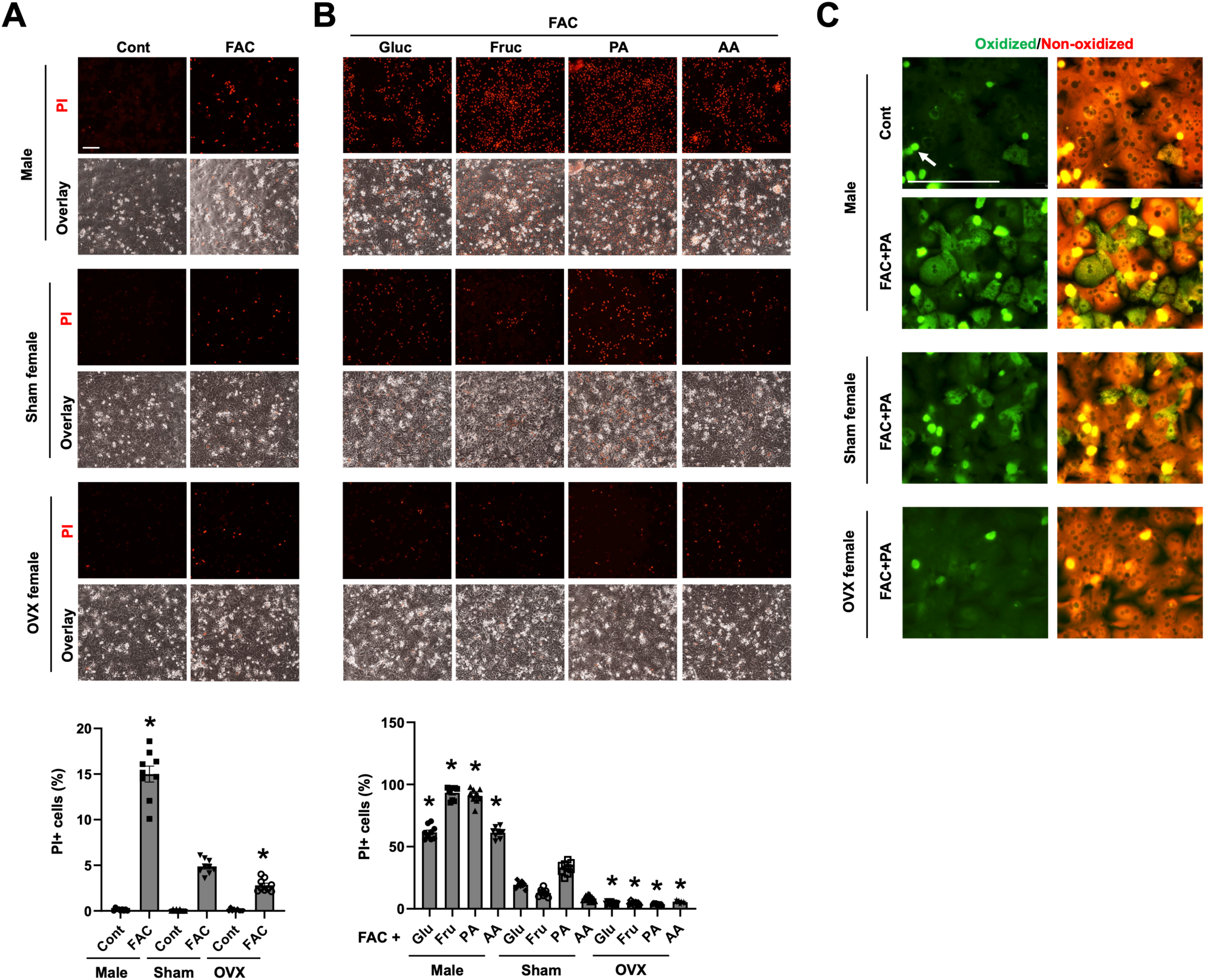
Ovariectomy enhances the resistance of female hepatocytes to ferroptosis. (**A**) P-Hepa of male, sham-female and OVX-female mice were treated with 50 μM FAC for 24 h, and ferroptotic cells was stained by PI. (**B**) P-Hepa were treated for 24 h by 50 μM FAC, along with 25 μM glucose (Glu), 25 μM fructose (Fru), 150 μM palmitic acid (PA), or 1x Amino Acids (AA), followed by PI staining. (**C**) P-Hepa were treated with or without the FAC and PA combination, and lipid peroxidation was determined by staining with BODIPY C11. Green fluorescence denotes oxidized lipids, and White arrow denotes dead cells that are present in all the presented images. The percentages of PI^+^ cells (**A** and **B**) are shown as mean ± SEM (n = 9 fields of 3 independent experiments). *, *P* < 0.01 compared to the corresponding treatment conditions in sham-female hepatocytes. Bars, 100 μm.

### Elevated FTH1 and FSP1 expression enhances the resistance of OVX hepatocytes to ferroptosis

To understand the mechanisms by which OVX potentiates hepatocyte resistance to ferroptosis, we first compared the expression of iron metabolism and antioxidant defense related genes in OVX and sham female hepatocytes. Interestingly, TfR1 mRNA expression further decreased by 37%, but FSP1 expression was increased by 100%, in OVX hepatocytes compared to the sham-female control (**Fig. 6A**). No significant changes were observed in the expression of other iron transporters and GPX4. Protein analysis further confirmed a reduction in TfR1 expression by 56% and an increase in FTH1 by 117% in OVX hepatocytes at the baseline (**Fig. 6B**). Following 6-h FAC treatment, the reduction of TfR1 expression was not significant between sham and OVX hepatocytes (**Fig. 6C**). In contrast, FTH1 expression was induced by 201% in sham hepatocytes and only by 67% in OVX hepatocytes (**Fig. 6C**). The lack of a difference in the reduction of TfR1, but in the induction of FTH1, mimics that between male and female hepatocytes in response to FAC (**Fig. 3E**).

**Figure 6.**
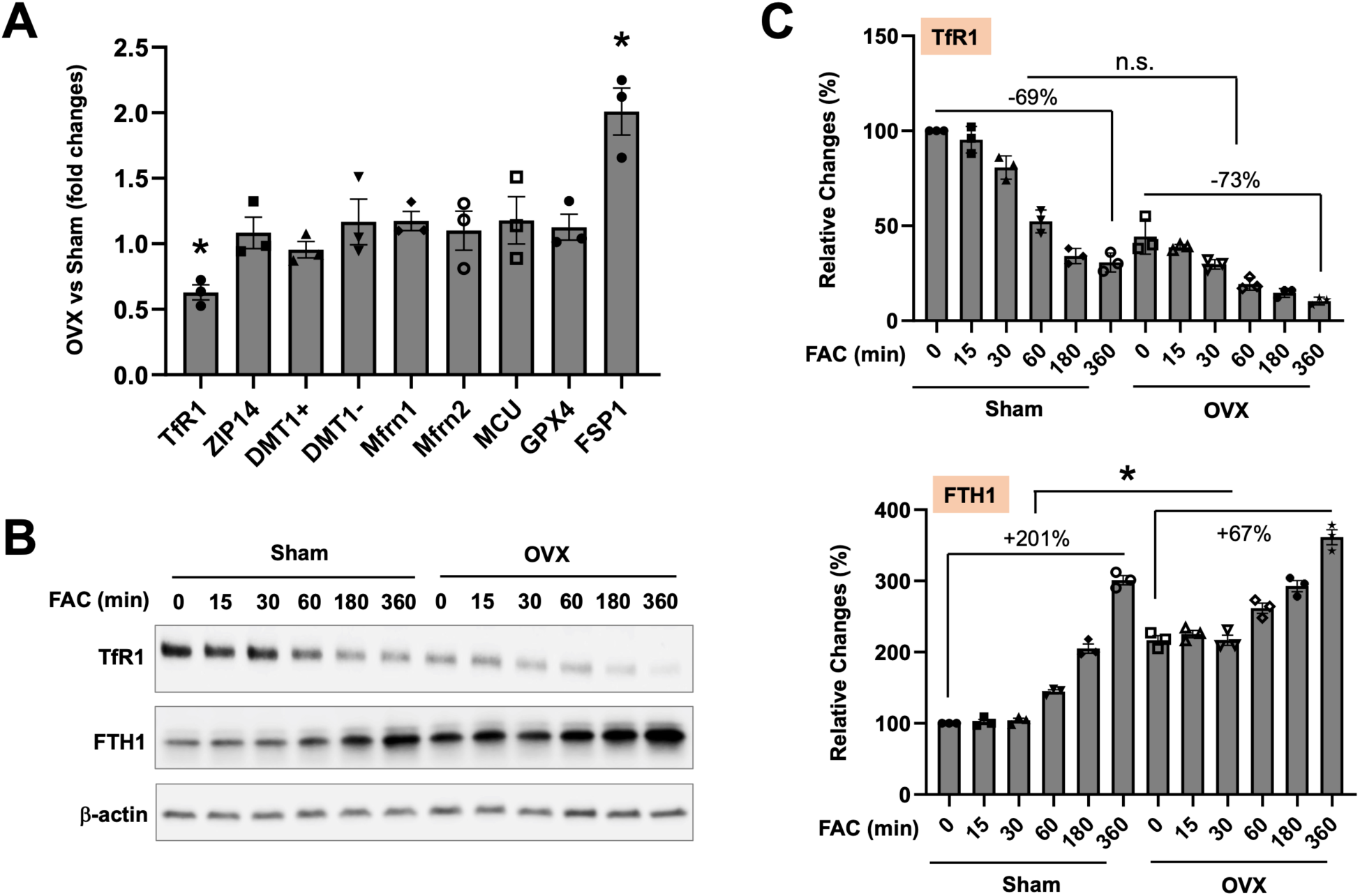
Ovariectomy alters the expression of TfR1, FTH1 and FSP1 expression in female hepatocytes. (**A**) The mRNA expression of iron metabolism and antioxidant genes was determined in P-Hepa of OVX *vs* sham-female mice. (**B**) P-Hepa of sham-female and OVX mice were treated with 100 μM FAC for the indicated times, and the protein expression of TfR1 and FTH1 was determined by Western blotting. (**C**) Quantified temporal changes in TfR1 and FTH1 expression are shown. Data are mean ± SEM (n = 3 independent experiments). *, *P* < 0.01.

Apart from the roles of TfR1 and FTH1, increased FSP1 expression can add another layer of protection. Indeed, inhibition of FSP1 with iFSP1 sensitized OVX hepatocytes to FAC-induced ferroptosis (**Fig. 7A**). To further determine the role and mechanism of elevated FSP1 expression in protecting hepatocytes, we generated HepG2 cells the expression of exogenous V5-tagged human FSP1 (V5-FSP1) (**Fig. 7B**). Compared to the control, HepG2/V5-FSP1 cells displayed significantly decreased expression of TfR1 and FTH1 by 72% and 41%, respectively. Mfrn1 expression was decreased by 33%. In response to FAC treatment, 4.8% backbone transduced control cells underwent ferroptosis. Strikingly, overexpression of FSP1 completely blocked FAC induced ferroptosis with only 0.3% cells positive for PI staining (**Fig. 7C**). HepG2/V5-FSP1 cells were also strongly resistant to RSL3 induced ferroptosis (**Fig. 7D**). We showed in the above that RSL3 increased mitochondrial Fe^2+^ loading and mtROS production. Intriguingly, FSP1 overexpression reduced the baseline and abolished RSL3 induced mtROS production and mitochondrial Fe^2+^ accumulation (**Fig. 7E**). The mechanism by which FSP1 suppresses mitochondrial iron loading is unknown, but it could be partially due to the lower expression of TfR1 and Mfrn1 (**Fig. 7B**). Our findings thus suggest that elevation in FSP1 expression inhibits hepatocyte ferroptosis through a previously unappreciated mechanism by preventing labile iron overload in the mitochondria.

**Figure 7.**
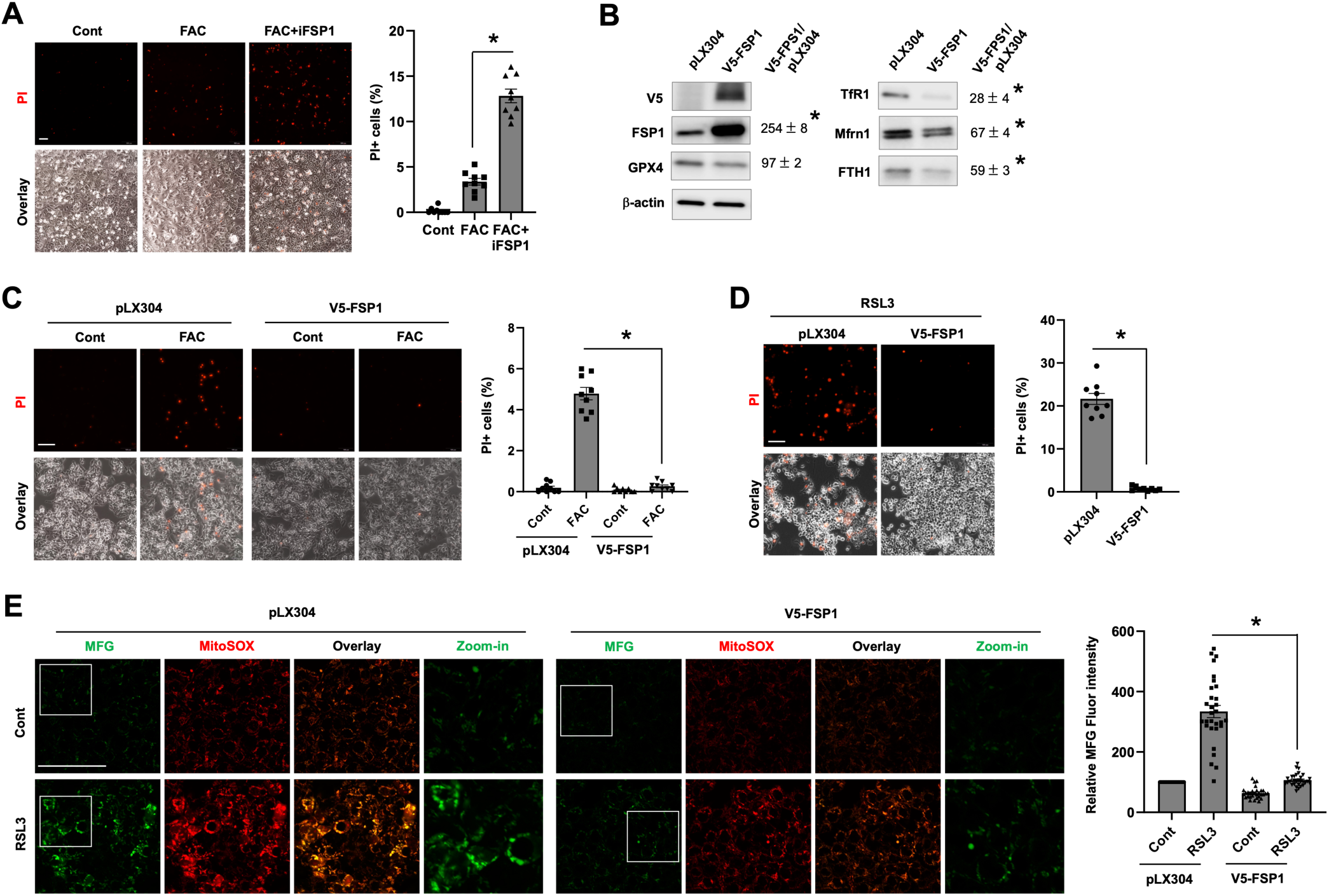
FSP1 protects hepatocytes against ferroptosis by attenuating mitochondrial iron loading and mtROS production. (**A**) OVX hepatocytes were treated for 24 h by 100 μM FAC in the presence or not of 10 μM iFSP1, and then stained by PI. (**B**) The effects of stable expression of V5-FSP1 on iron metabolism and antioxidant defense genes in HepG2 cells. (**C**) HepG2 cells with the stable expression of V5-FSP1 or the backbone control (pLX304) were treated with or not 100 μM FAC for 24 h, followed by PI staining. (**D**) HepG2/V5-FSP1 and control cells were treated for 4 h with 200 nM RSL3 prior to PI staining. The percentages of PI^+^ cells (**A**, **C** and **D**) were quantified, and the data shown are mean ± SEM (n = 9 fields of 3 independent experiments). (**E**) HepG2/V5-FSP1 and control cells were treated for 4 h with 200 nM RSL3, followed by staining for mitochondrial Fe^2+^ with 5 μM Mito-FerroGreen (MFG) and mtROS with 5 μM MitoSOX. The fluorescence intensity of MFG staining was quantified and is presented as mean ± SEM (n = 30 cells of 3 independent experiments). *, *P* < 0.01. Bars, 100 μm.

### OVX induced hyperactivation of ERK protects hepatocytes against ferroptosis

A plethora of cellular signaling regulates cell survival including MAPK (ERK1/2 and p38 MAPK) and Akt mediated signaling [39, 40]. We identified that FAC treatment for 4 h resulted in a significant increase in the phosphorylation of ERK1/2 and Akt in the more responsive male hepatocytes. The increase in phospho-ERK1/2 was mitigated or completed blocked by MAP kinase kinase (MEK) inhibitor U0126, calcium blocker BAPTA-AM, and lipid radical scavenger liproxstatin-1 (Lip-1). The upregulation of phospho-Akt was only prevented by Lip-1 (**Fig. S3A**). It is suspected that iron-induced activation of ERK and Akt is a feedback protective mechanism against lipid peroxidation-mediated cellular damage. Indeed, inhibition of ERK or Akt prior to FAC treatment exacerbated FAC-induced ferroptosis as indicated by the significantly increased numbers of PI^+^ cells (**Fig. S3B**).

It was previously shown that OVX increases the phosphorylation/activity of p38 MAPK in rat cardiomyocytes [41]. We wondered whether OVX induces the phosphorylation of ERK and Akt in hepatocytes. Intriguingly, the baseline phospho-ERK1/2 level was markedly higher in OVX hepatocytes than the sham control. Phospho-Akt was only slightly increased in OVX hepatocytes (**Fig. 8A**). The potential roles of ERK and Akt in mediating the resistance of OVX hepatocytes to iron-induced ferroptosis was assessed by using PD0325901, a MEK inhibitor that blocks ERK activation, and MK2206, a selective Akt inhibitor. We showed that inhibition of ERK and Akt both increased the susceptibility of OVX hepatocytes to FAC-induced ferroptosis (**Fig. 8B**). AML12 cells are transgenic mouse hepatocytes the overexpression of TGF-α, which is a potent agonist of ERK signaling [42]. We have previously shown that, compared to P-Hepa and HepG2 cells, AML12 cells are resistant to FAC-induced ferroptosis [18]. Indeed, phosphorylation of ERK1/2 was much stronger in AML12 cells relative to P-Hepa (**Fig. 8C**), and inhibition of ERK by PD0325901 sensitized AML12 cells to FAC-induced ferroptosis (**Fig. 8D**). A recent study showed that MAPK potentially upregulates FSP1 expression in KRAS-mutant cancer cells [43]. By using the AML12 cell model we could show that inhibition of ERK significantly reduced the mRNA expression of FSP1 (**Fig. 8E**). The subcellular localization of V5-FSP1 in HepG2 cells did not seem significantly altered by the inhibition of ERK (**Fig. S4**). These results suggest that OVX induces the hyperactivation of ERK, which contributes to increased defense of OVX hepatocytes against ferroptosis in part through upregulation of FSP1.

**Figure 8.**
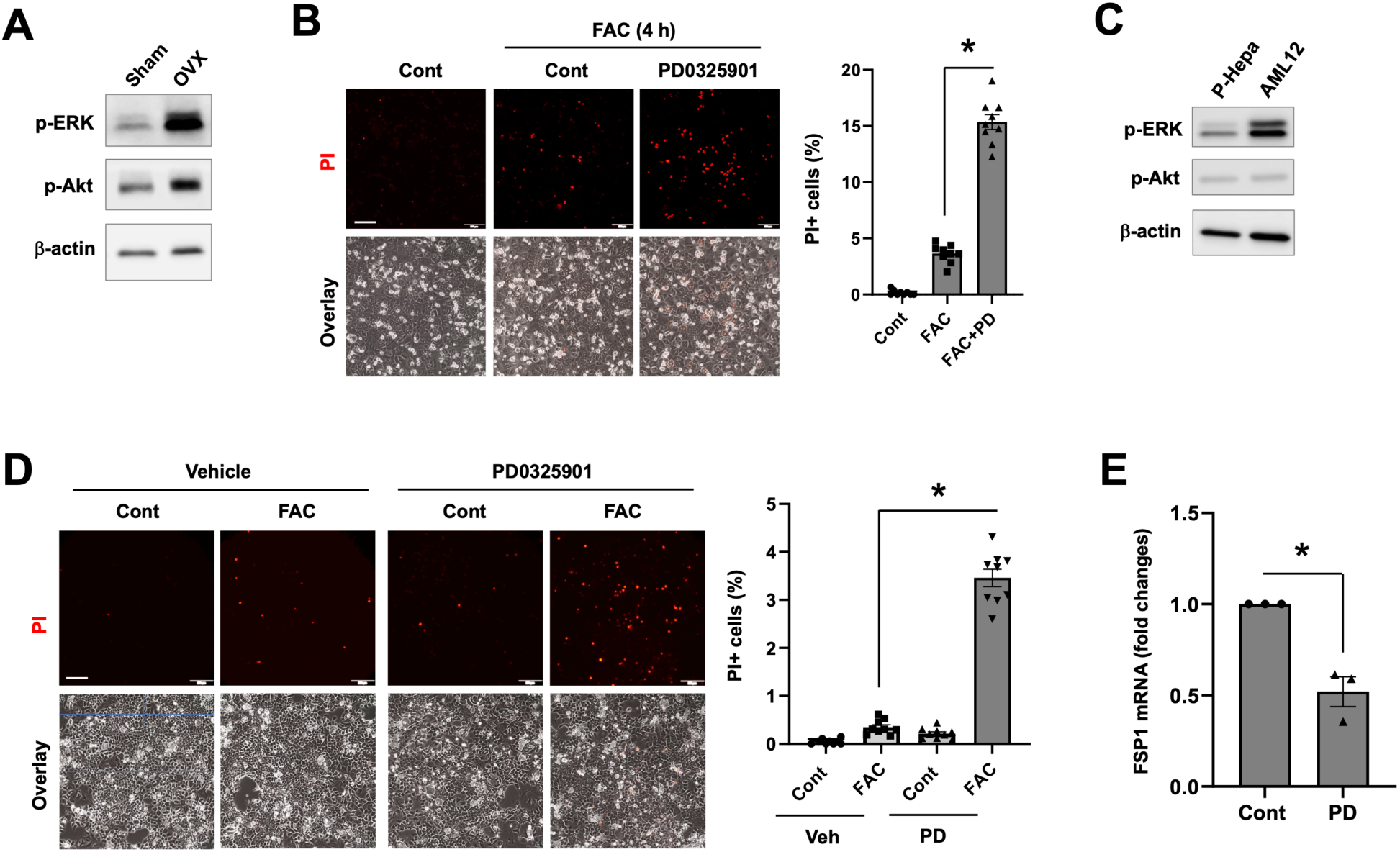
Ovariectomy induces hyperactivation of ERK protecting hepatocytes against ferroptosis. (**A**) The expression levels of phospho-ERK1/2 (p-ERK) and phospho-Akt (p-Akt) were determined in P- Hepa of OVX and sham-female mice. (**B**) OVX hepatocytes were treated for 4 h with 100 μM FAC in the presence or not of 5 μM PD0325901 (PD), and then stained for PI. (**C**) The expression of p-ERK and p- Akt in male P-Hepa and AML12 cells. (**D**) AML12 cells were treated for 24 h with or without FAC or PD, followed by PI staining. The percentages of PI^+^ cells (**B** and **D**) were quantified and are shown as mean ± SEM (n = 9 fields of 3 independent experiments). (**E**) AML12 cells were treated for 4 h with or without PD, and the mRNA expression of FSP1 was determined by qRT-PCR. Data are mean ± SEM (n = 3). *, *P* < 0.01. Bars, 200 μm.

## Discussion

Ferroptosis, an iron-catalyzed lipid peroxidation-dependent oxidative death, has been increasingly recognized as an important mechanism in the pathogenesis and progression of many diseases including the liver diseases [44, 45], which display sexual dimorphism. Our study for the first time showed that male and female hepatocytes manifest differential vulnerability to ferroptosis induction by various inducers including excess iron and pharmacological drugs. A lower susceptibility of female hepatocytes to ferroptosis is in part due to the higher expression of FTH1 and lower expression of TfR1 and Mfrn1 at the baseline (**Fig. 9**). The resistance of female hepatocytes to ferroptosis is not attributable to female hormone mediated signaling as hepatocytes derived from OVX mice displayed increased resistance to ferroptosis. Elevation in ferroptosis resistance of the OVX hepatocyte attributes to multiple mechanisms including decreased TfR1 expression, upregulated expression of FTH1 and FSP1, and hyperactivated ERK signaling (**Fig. 9**).

**Figure 9.**
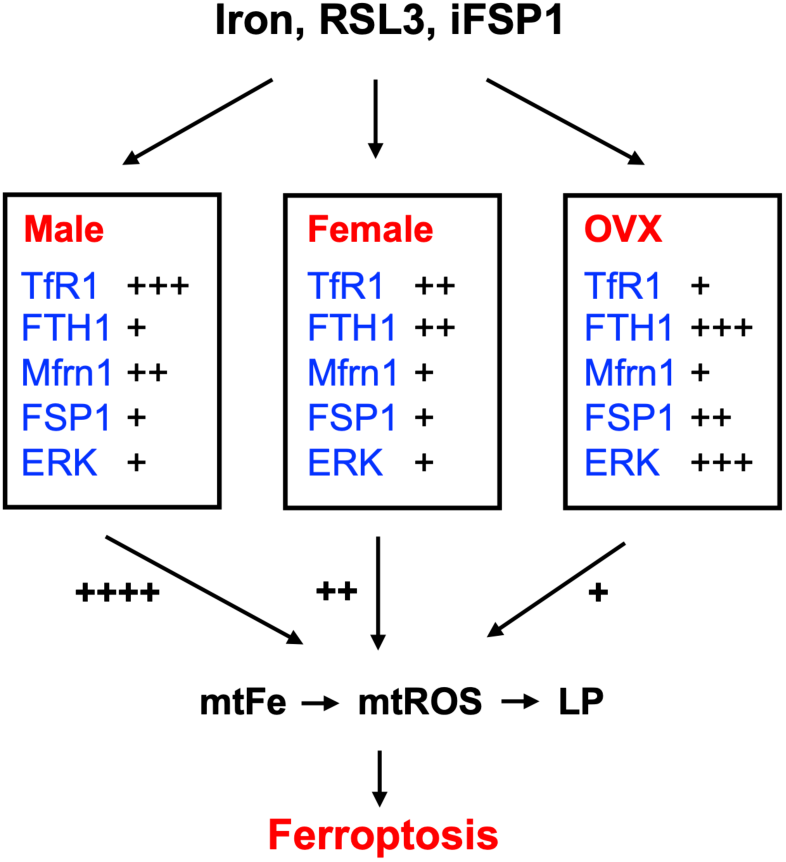
A schematic model demonstrating the differential susceptibility of male, female and OVX hepatocytes to induced ferroptosis. The contents inside the boxes show the expression levels (TfR1, FTH1, Mfrn1 and FSP1) or activity (ERK) of proteins that determine the sensitivity of hepatocytes to ferroptosis inducers including iron excess, RSL3, and iFSP1. The listed factors regulate mtFe content, mtROS production capacity and the magnitude of LP that leads to ferroptosis. *mtFe*, mitochondrial Fe^2+^; and *LP*, lipid peroxidation.

Hepatocytes in stressed conditions can undergo different types of death including apoptosis, necrosis, and ferroptosis [46]. Sex differences in hepatic death have previously been demonstrated. It was shown that female mouse hepatocytes are more susceptible to ethanol-induced apoptosis than male hepatocytes [47]. In contrast, Gupta *et al*. showed that herbicide diquat induces a higher rate of hepatic necrosis and more severe liver damage in male rats [48]. Acetaminophen overdose induced acute liver injury is more severe in male mice [49], but the type of hepatic death is unclear. Earlier reports showed that acetaminophen overdose induces necrosis of the hepatocytes [50], while multiple recent studies have identified that acetaminophen causes hepatic ferroptosis [51, 52]. Our current study by using *in vitro* model demonstrated that male hepatocytes are more susceptible to ferroptosis than female hepatocytes. Our finding is consistent with a recent report that renal proximal tubular epithelial cells of male mice are more vulnerable to ferroptosis by utilizing the GPX4 knockout mouse model [17]. That study showed that differential NRF2 expression is the underlying cause of the gender difference in renal epithelial ferroptosis. Our work failed to identify NRF2 as a candidate mechanism for the sexual dimorphism in hepatocyte ferroptosis. Instead, we showed that gender difference in hepatocyte ferroptosis by excess iron, RSL3 and iFSP1 is associated with distinct mitochondrial iron metabolism and mtROS production, at least in mice. Previous evidence showed that RSL3 induces ferroptosis by inactivating GPX4 [33], and iFSP1 acts by inhibiting FSP1-mediated production of CoQH_2_ [35]. To the best of our knowledge, our study for the first time identified that RSL3 and iFSP1 induced ferroptosis is also in part through the induction of mitochondrial iron overload.

The molecular mechanisms underlying differential mitochondrial iron metabolism between male and female hepatocytes are multifold. We identified differential expression of TfR1, FTH1, and Mfrn1 in the hepatocytes of male and female mice and/or humans. TfR1 is a ubiquitous iron transporter that takes up transferrin-bound iron from the circulation or other external sources. It is conceivable that a higher basal level of TfR1 leads to a more robust intracellular iron loading than cells that bear less TfR1. However, we failed to find a difference in the rate of reduction in TfR1 expression in male, female and OVX hepatocytes in response to excessive external iron. This could be explained by that less labile iron than the-actually-loaded amount is required to trigger the response or degradation of TfR1. The correlation of TfR1 expression with the rate of ferroptosis is well noted [37]. By contrast, the role of FTH1 in ferroptosis is a paradox. On the one side, deficiency in FTH1, a major iron storage protein, potentially leads to labile Fe^2+^ accumulation, which could induce lipid peroxidation and ferroptosis. On the other hand, high intracellular FTH1 expression, which indicates abundant iron storage, could lead to the release of more free iron through ferritinophagy [53]. Our current results suggest that mitochondrial integrity is impaired by FTH1 deficiency as indicated by mtROS accumulation. The higher baseline mtROS is probably the primary mechanism underlying the high vulnerability of FTH1-deficient cells to FAC and RSL3 induced ferroptosis. This speculation is supported by our previous report showing that P-Hepa pretreated with rotenone [54], an inhibitor of mitochondrial complex I that increases mtROS content, were highly sensitive to FAC-induced ferroptosis [18]. Our finding is further supported by a previous report that hepatocytes with the knockout of FTH1 are highly susceptible to liver damage in mice fed a high-iron diet [55]. Mfrn1 is an inner mitochondrial membrane protein that mediates iron import into the matrix of the mitochondria. Mfrn1 expression is particularly important in erythroid precursors for heme production [56]. However, knockout of Mfrn1 in hepatocytes does not seem to significantly alter mitochondrial iron homeostasis in the physiological state [57]. Therefore, despite the important role of mitochondrial iron loading in ferroptosis, the role of Mfrn1 in regulating ferroptosis response remains unclear. Our current data demonstrated that HepG2 cells with lowered Mfrn1 expression displayed decreased sensitivity to FAC-induced ferroptosis. However, whether the reduced sensitivity is due to a direct role of Mfrn1 deficiency or indirectly via downregulated TfR1 expression remains not understood. We cannot completely exclude a potential role of antioxidant protection in sexual dimorphism of hepatocyte ferroptosis despite the lack of differential expression in NRF2, GPX4 and FSP1 between male and female hepatocytes. The abundance of glutathione and CoQ may be examined in future studies, and yet unidentified antioxidant defense mechanisms may play a role.

Another key finding of the current study is that OVX promotes the resistance of female hepatocytes to ferroptosis. We have identified that OVX induced decreased TfR1 and increased FTH1 expression as the mechanisms of elevated resistance of OVX hepatocytes to ferroptosis. This finding is consistent with a previous finding in the adipose tissues of OVX rats, whereby OVX achieved the effects by decreasing RNA binding activity of iron regulatory protein 1 (IRP1) [58]. We also demonstrated for the first time that FSP1 is significantly increased in OVX hepatocytes. Elevation in FSP1 expression provided strong protection against both iron and RSL3 induced ferroptosis. Previous evidence suggests that FSP1 inhibits ferroptosis by promoting the production of CoQH_2_ at the plasma membrane [16]. We have novel finding that increasing FSP1 expression mitigated RSL3-induced mitochondrial Fe^2+^ loading and mtROS production, thereby suggesting a previously unappreciated role of FSP1 in protecting mitochondrial integrity. The mechanism underlying the upregulated expression of FSP1 in OVX hepatocytes is unknown but could be due to the hyperactive ERK signaling because inhibition of ERK attenuated FSP expression in AML12 cells, a cell line that mimics OVX hepatocytes in terms of ERK hyperactivation. Our finding is consistent with a recent report that increased expression of FSP1 in KRAS-mutant cancer cells is associated with mitogen-activated protein kinase (MAPK) and NRF2 [43]. Moreover, our finding of the hyperactivation of ERK in OVX hepatocytes provides a novel mechanistic insight into upregulated hepcidin production in women of post-menopause, because MAPK is known to mediate the activation of hepcidin expression [59, 60].

In conclusion, our study has demonstrated that gender difference in hepatocellular iron handling particularly in mitochondrial iron metabolism underlies sexual dimorphism in the vulnerability of hepatocytes to ferroptosis. The study was primarily based on the use of *in vitro* cultured hepatocytes, but it is plausible to speculate that sexual dimorphism in hepatic ferroptosis should also exist in *in vivo* given the similar changes in TfR1, Mfrn1 and FTH1 expression between the liver and cultured hepatocytes.

## Acknowledgements

This study was supported by National Institute of Health grant R01DK125647. We would like to thank Dr. Adam Gracz for sharing the Olympus IX83 live imaging microscope.

## Author contributions

Conception and design: PH; Acquisition of data: HT, HYD, CT, SB, ESG, PH; Analysis and interpretation of data: SS, LZ, RP, PH; Writing, review, and revision of the manuscript: ESG, SS, LZ, RP, PH.

## Abbreviations

ALD: alcoholic liver disease
CoQH_2_: reduced coenzyme Q10
DMT1: divalent metal transporter 1
FAC: ferric ammonium citrate
FSP1: ferroptosis suppressor protein 1
FTH1: ferritin heavy chain 1
GPX4: glutathione peroxidase 4
IRE: iron-responsive element
MCU: mitochondrial Ca^2+^ uniporter
mtROS: mitochondrial ROS
NAFLD: non-alcoholic fatty liver disease
NRF2: nuclear factor-erythroid factor 2- related factor 2
NTBI: non-transferrin-bound iron
OVX: ovariectomy
P-Hepa: primary mouse hepatocytes
PI: propidium iodide
TfR1: transferrin receptor 1
TGF-α: transforming growth factor α
ZIP14: Zrt- and Irt-like (ZIP) protein 14

