## Supplemental data for "Differences in Hepatocellular Iron Metabolism Underlie Sexual Dimorphism in Hepatocyte Ferroptosis"

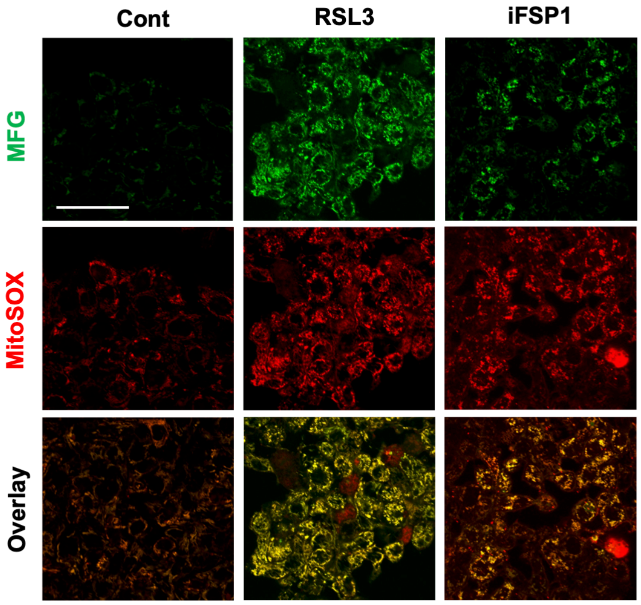


**Figure S1. The effects of RSL3 and iFSP1 on mitochondrial iron loading and mtROS production in HepG2 cells.** HepG2 cells were treated for 4 h with 200 nM RSL3, 24 h with 10 μM iFSP1, or the vehicle. Mitochondrial $Fe^{2+}$ was then labelled by Mito-FerroGreen and mtROS by MitoSOX. Bar, 50 μm.

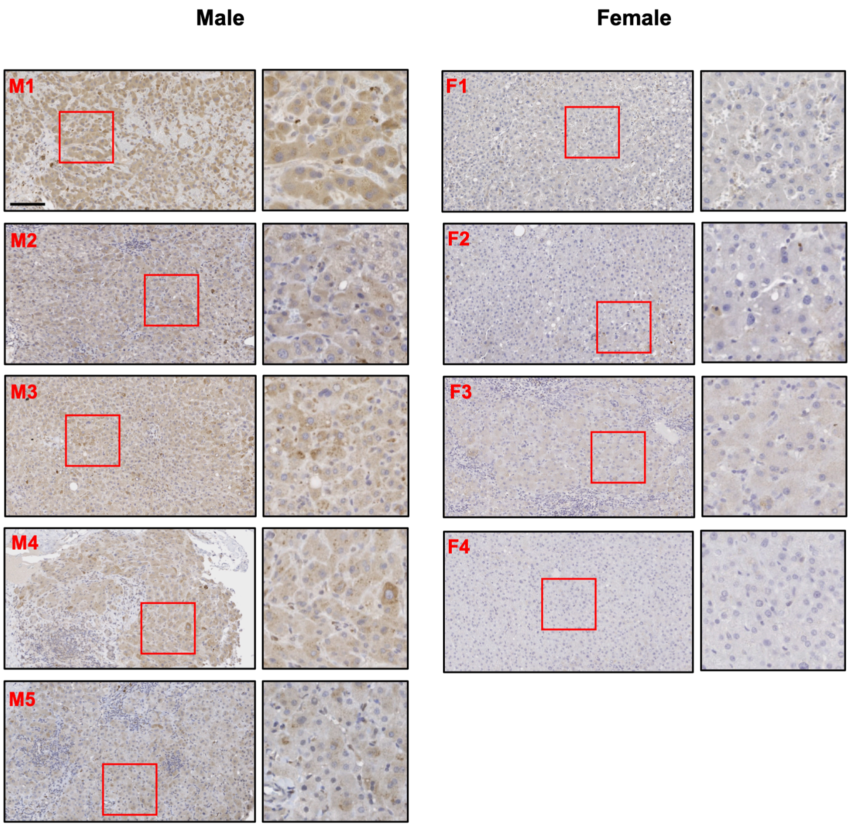


**Figure S2. Representative immunohistochemical images showing the staining of Mfrn1 in the liver of male and female human subjects.** *M*, male; and *F*, female. Bar, 100 μm.

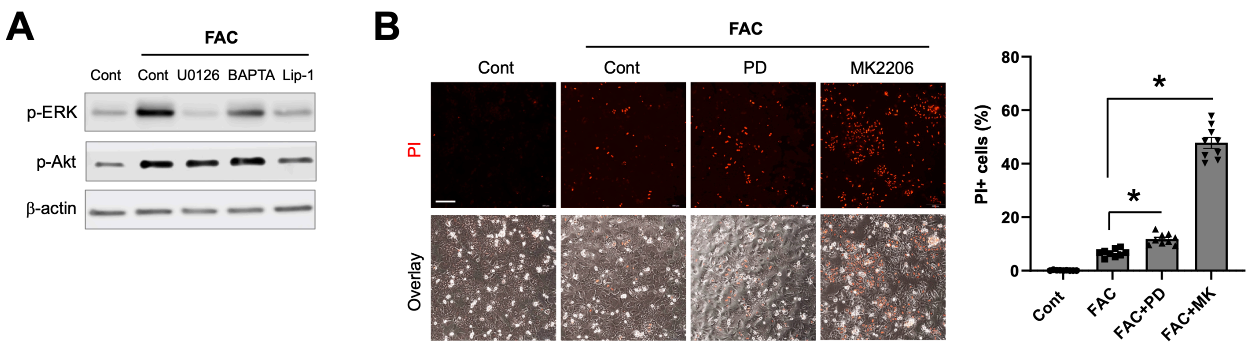


**Figure S3. ERK and Akt protect against hepatocyte ferroptosis.** (**A**) The expression of p-ERK and p-Akt in male P-Hepa that were treated for 3 h with 100 μM FAC in the presence of 10 μM U0126 (a MEK inhibitor), 40 μM BAPTA-AM (BAPTA, a calcium chelator), 5 μM liproxstatin-1 (Lip-1, a lipid radical scavenger), or vehicle. (**B**) PI staining of male P-Hepa that were treated for 4 h with 20 μM FAC in the presence of 5 μM PD (a MEK inhibitor), 5 μM MK2206 (MK, an Akt inhibitor), or vehicle. The percentages of $PI^+$ cells are shown as mean ± SEM (n = 9 fields of 3 independent experiments). *, *P* < 0.01. Bar, 200 μm.

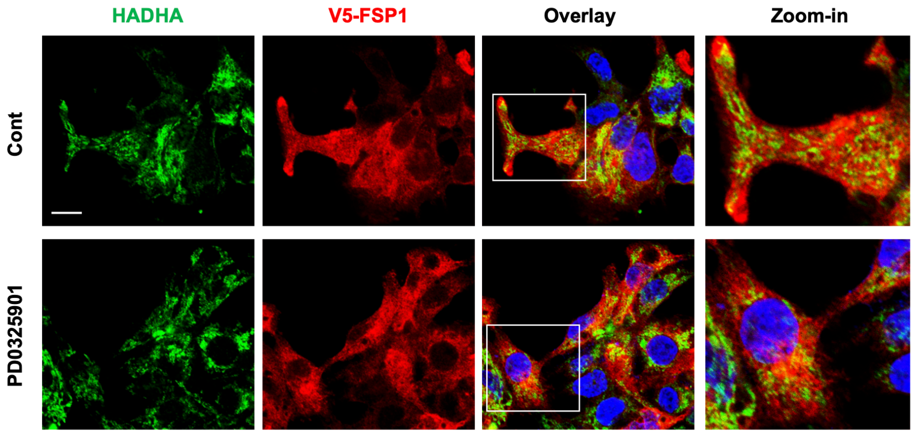


**Figure S4. The effect of ERK inhibition on subcellular localization of FSP1.** HepG2/V5-FSP1 cells were treated with or without 5 μM PD0325901, followed by co-staining with anti-V5 and anti-HADHA (an inner mitochondrial membrane protein) antibodies and confocal imaging. Bar, 10 μm.

**Supplementary Table 1. List of reagents and antibodies**

| Name | Source | Catalog | Species |
|---|---|---|---|
| **Reagents** | | | |
| (1S, 3R)-RSL3 | Cayman | 19288 | |
| BAPTA-AM | Sigma-Aldrich | A1076 | |
| Bicinchoninic Acid Assay kit | Sigma-Aldrich | BCA1 | |
| Blasticidin | Invitrogen | R210-01 | |
| Cell Lysis Buffer (10x) | Cell Signaling Technology | 9803 | |
| cOmplete Phosphatase Inhibitor | Roche | 04906837001 | |
| cOmplete Protease Inhibitor (EDTA free) | Roche | 11836170001 | |
| Dexamethasone | Sigma-Aldrich | D2915 | |
| D-(+)-Glucose | Sigma-Aldrich | G7021 | |
| D-(-)-Fructose | Sigma-Aldrich | F3510 | |
| DMEM/F12 | Gibco | 21331-020 | |
| Ferric Ammonium Citrate | Sigma-Aldrich | F5879 | |
| Fetal Bovine Serum | R&D Systems | S11150 | |
| FIN56 | Cayman | 25180 | |
| HBSS (10x) | Gibco | 14185-052 | |
| HEPES (1M) | Gibco | 15630-080 | |
| High-Capacity cDNA Reverse Transcription Kit | Applied Biosystems | 43-688-13 | |
| iFSP1 | Cayman | 29483 | |
| Image-iT Lipid Peroxidation Kit | Thermo Fisher Scientific | C10445 | |
| Insulin | Sigma-Aldrich | I0516 | |
| Insulin-Transferrin-Selenium | Gibco | E2311 | |
| Liberase | Sigma-Aldrich | 05401119001 | |
| Liproxstatin-1 | Cayman | 17730 | |
| MEM Amino Acids (50x) | Thermo Fisher Scientific | 11130-051 | |
| Mito-FerroGreen | Dojindo | M489-10 | |
| MitoSOX Red | Thermo Fisher Scientific | M36008 | |
| MK2206 | Cayman | 11593 | |
| Palmitic acid | Sigma-Aldrich | P5585 | |
| Paraformaldehyde | Sigma-Aldrich | P6148 | |
| PD0325901 | Cayman | 13034 | |
| Penicillin/Streptomycin (P/S) | Thermo Fisher Scientific | 15140122 | |
| Percoll (40%) | Sigma-Aldrich | P1644 | |
| Polybrene | Sigma-Aldrich | TR-1003 | |
| *Power* SRBR Green PCR Master Mix | Applied Biosystems | 43-676-59 | |
| Prolong Diamond Antifade Mounting Medium | Thermo Fisher Scientific | P36961 | |
| Propidium Iodide (PI) | Invitrogen | P1304MP | |
| Puromycin | Sigma-Aldrich | P8833 | |
| RNeasy Mini Kit | Qiagen | 74136 | |
| RPMI Medium | Sigma-Aldrich | R8758 | |

| | | | |
|---|---|---|---|
| Trypsin-EDTA (0.25%) | Thermo Fisher Scientific | 15400-054 | |
| U0126 | Cayman | 70970 | |
| William’s E Medium | Sigma-Aldrich | W1878 | |
| **Antibodies** | | | |
| anti-mouse Alexa Fluor 488 | Thermo Fisher Scientific | A11031 | Goat |
| anti-mouse HRP | Cell Signaling Technology | 7076 | Goat |
| anti-rabbit Alexa Fluor 568 | Thermo Fisher Scientific | A11034 | Goat |
| anti-rabbit HRP | Cell Signaling Technology | 7074 | Goat |
| β-actin | Sigma-Aldrich | A2066 | Rabbit |
| FTH1 | Cell Signaling Technology | 3998 | Rabbit |
| FSP1 | Cell Signaling Technology | 24972 | Rabbit |
| GPX4 | Cell Signaling Technology | 59735 | Rabbit |
| HADHA | Santa Cruz Biotechnology | sc-374497 | Mouse |
| Mitoferrin 1 | Proteintech | 26469-1-AP | Rabbit |
| NRF2 | Cell Signaling Technology | 12721 | Rabbit |
| Phospho-ERK1/2 | Cell Signaling Technology | 3179 | Rabbit |
| Phospho-Akt | Thermo Fisher Scientific | PA5-102853 | Rabbit |
| TfR1 | Thermo Fisher Scientific | 13-6800 | Mouse |
| V5 | Cell Signaling Technology | 13202 | Rabbit |

**Supplementary Table 2. Information of male (M) and female (F) human subjects**

| **Patient** | **Age** | **Diagnosis** |
|---|---|---|
| M1 | 62 | Colorectal cancer liver metastasis |
| M2 | 57 | Colorectal cancer liver metastasis |
| M3 | 54 | Cholangiocarcinoma |
| M4 | 78 | Cholangiocarcinoma |
| M5 | 76 | Cholangiocarcinoma |
| F1 | 58 | Colorectal cancer liver metastasis |
| F2 | 63 | Colorectal cancer liver metastasis |
| F3 | 59 | Cholangiocarcinoma |
| F4 | 48 | Colorectal cancer liver metastasis |

**Supplementary Table 3. List of primers and sequences**

| Gene | Sequence |
|---|---|
| mDMT1-IRE F | TGGGTCTGTCTTTCCTGGAC |
| mDMT1-IRE R | TGCAACAGCACATACTTGTGG |
| mDMT1-non-IRE F | TGGGTCTGTCTTTCCTGGAC |
| mDMT1-non-IRE R | GAACAAGCTCACCTCCGAAC |
| mFSP1 F | CGATACCAAGGAGCCCAAGA |
| mFSP1 R | TAGAGTTCTCACATGGCCGG |
| mGPX4 F | CGATCTGCATGCCCGATATG |
| mGPX4 R | GGCATCGTCCCCATTTACAC |
| mMCU F | TGCCAATTCACACTCAAGCC |
| mMCU R | AAGTCATCGAGGAGCAGGAG |
| mSLC25A28 F | CTGGACACATCACAGGCA |
| mSLC25A28 R | ATTCCTCTTGCCGCTTTGTG |
| mSLC25A37 F | TCTCACAACCATCACCCTCC |
| mSLC25A37 R | AGCAACACTAGGCCACTCTT |
| mTfR1 F | AATGGGTGTGTTGAAGACAA |
| mTfR1 R | ACATTCTCAGGTGGCAGCTT |
| mZIP14 F | CTGGCTATTGGTGCCTCCTTCA |
| mZIP14 R | TGCCAGCATTGAGCAGGATGAC |
| mβ-actin F | AGCCATGTACGTAGCCATCC |
| mβ-actin R | TCTCAGCTGTGGTGGTGAAG |

Note: m, mouse.
